# Optimal Frequency for Seizure Induction with Electroconvulsive Therapy and Magnetic Seizure Therapy

**DOI:** 10.1101/2024.09.28.615333

**Authors:** Angel V. Peterchev, Zhi-De Deng, Christopher Sikes-Keilp, Elyssa C. Feuer, Moacyr A. Rosa, Sarah H. Lisanby

**Affiliations:** Department of Psychiatry and Behavioral Sciences, Duke University, Durham, NC, USA; Department of Biomedical Engineering, Duke University, Durham, NC, USA; Department of Electrical and Computer Engineering, Duke University, Durham, NC, USA; Department of Neurosurgery, Duke University, Durham, NC, USA; Noninvasive Neuromodulation Unit, Experimental Therapeutics and Pathophysiology Branch, Intramural Research Program, National Institute of Mental Health, National Institutes of Health, Bethesda, MD, USA; Department of Psychiatry, University of North Carolina, Chapel Hill, NC, USA; University of Toledo College of Medicine and Life Sciences, Toledo, OH, USA; Institute for Advanced Research in Neurostimulation, São Paulo, SP, Brazil

## Abstract

Electroconvulsive therapy (ECT) and magnetic seizure therapy (MST) are effective in the treatment of medication-resistant depression. Determining the stimulus frequency resulting in the lowest seizure threshold could produce fewer adverse effects by reducing the overall stimulus intensity. To determine the optimal frequency for seizure induction, four male rhesus macaques were titrated with an increasing number of pulses at fixed frequencies ranging from 5 to 240 pulses per second (pps) using ultrabrief-pulse right-unilateral ECT and circular-coil-on-vertex MST. The seizure threshold dependence on stimulus frequency was similar for ECT and MST. While higher frequencies required progressively shorter trains to induce a seizure, the middle frequency range was associated with the fewest pulses (and hence the least charge and energy), with a minimum at 16 pps and similarly low thresholds for 10 and 25 pps. The number of pulses at seizure threshold increased markedly at lower and higher frequencies. The lowest stimulus frequencies, 5 and 10 pps, were associated with the greatest ictal power measured by electroencephalography. While this study did not assess efficacy or side effects, the results highlight the significance of stimulus frequency for seizure induction, suggest efficient titration schedules that minimize exposure to the electrical stimulus, and can inform studies to assess the impact on clinical outcomes.

## Introduction

Electroconvulsive therapy (ECT) is highly effective for medication-resistant depression (APA, 2001a). Manipulations of stimulus parameters, such as the pulse width, electrode placement, and number of pulses (or, equivalently, total charge), can significantly reduce side effects while maintaining efficacy (Peterchev *et al*, 2010b; Sackeim *et al*, 1993; Sackeim *et al*, 2000; Sackeim *et al*, 2008). Magnetic seizure therapy (MST) (Lisanby *et al*, 2001) can achieve similar antidepressant effects (Deng *et al*, 2024; Fitzgerald *et al*, 2018; Kayser *et al*, 2011; Kayser *et al*, 2015) but with a potentially more favorable side effect profile than ECT (Deng *et al*, 2024; Lisanby *et al*, 2003a; Moscrip *et al*, 2006; Polster *et al*, 2015; Spellman *et al*, 2008), ostensibly due to its ability to induce seizures with less electric field exposure (Lee *et al*, 2016). Many of the parameters associated with seizure therapies have yet to be examined for their clinical impact.

Pulse train frequency^1^ is commonly manipulated to individualize dose (APA, 2001a), but its effects on seizure threshold (ST) and clinical outcomes are poorly understood. Literature suggests that low frequencies are superior for seizure induction (Peterchev *et al*, 2010b). Stimuli in the range of 3–20 pps are optimal for photic seizure induction (Bickford *et al*, 1952) and stimuli in the range of 25–60 pps are optimal for the induction of after-discharges, considered the precursors of seizure (Swinyard, 1972). Indeed, lower frequencies (20–60 pps) appear to be more efficient than higher frequencies for seizure induction in clinical ECT and MST (Daskalakis *et al*, 2020; Girish *et al*, 2003; Swartz and Larson, 1989; Weaver *et al*, 1982), and titrating dose by increasing train duration results in lower ST than increasing frequency (Devanand *et al*, 1998). However, the most efficient stimulus frequencies for seizure induction have not been mapped.

Here we report the dependence of ST on stimulus frequency over a wide frequency range in nonhuman primates, and show that there is a narrow frequency band that minimizes the ST in both ECT and MST.

## Materials and Methods

This study was approved by the Institutional Animal Care and Use Committees of New York State Psychiatric Institute, Columbia University, and Duke University. We have previously reported detailed methods for nonhuman primate models of ECT and MST (Lisanby *et al*, 2003b; Moscrip *et al*, 2004).

### Subjects

Subjects were four male pathogen-free rhesus macaques (*Macaca mulatta*). Age and weights are given in Table 1.

**Table 1.** Age and weight range and individualized stimulation current amplitude used for all procedures in the four nonhuman primate subjects. The ECT and MST current amplitudes are given in milliamperes (mA) and percent of maximum stimulator output (% MSO), respectively.

|  |  | Age range<br>(years) | Weight range<br>(kg) | Stimulation amplitude |  |
| --- | --- | --- | --- | --- | --- |
|  |  |  |  | ECT (mA) | MST (% MSO) |
| Subject | CH | 11.1–13.6 | 8.5–9.9 | 565 | 86 |
|  | DY | 9.9–12.4 | 10.7–12.8 | 380 | 76 |
|  | MA | 10.1–12.3 | 8.3–9.8 | 230 | 66 |
|  | RZ | 17.2–18.8 | 8.1–12.1 | 280 | 84 |

### Study design

Two modalities were studied: ECT with right unilateral (RUL) electrode placement and MST with circular coil on vertex configuration, illustrated in Figure 1. ST was determined for a range of stimulus pulse train frequencies. One modality × frequency condition was tested per session. Each condition was repeated 3 times per subject. Sessions were separated by at least 5 days to minimize carryover effects.

**Figure 1.**
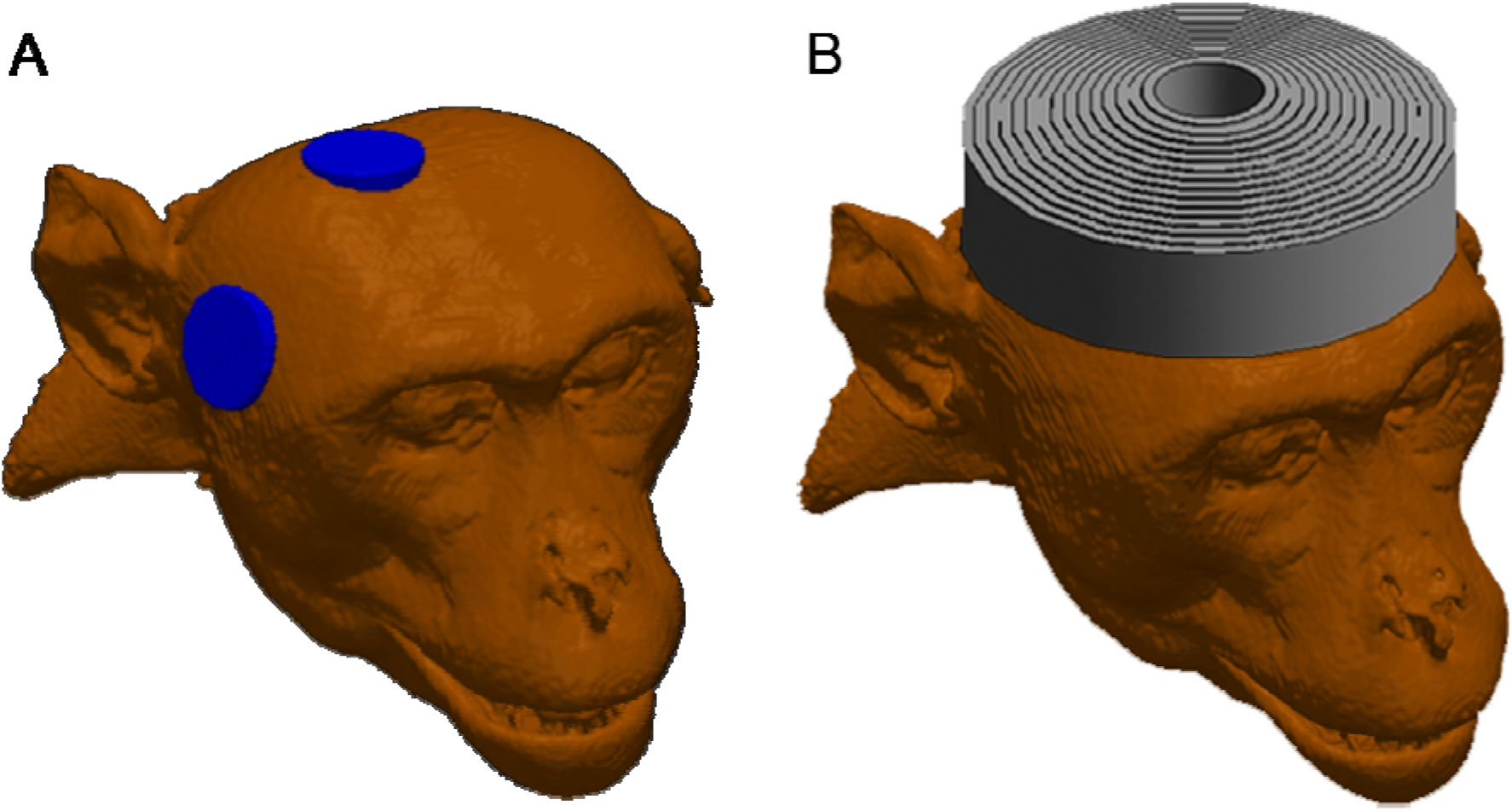
Illustration of the stimulus delivery configurations in a representative nonhuman primate subject. A: Right unilateral (RUL) electrode ECT configuration. B: Circular coil on vertex MST configuration.

### Anesthesia and monitoring

Anesthesia and monitoring followed previously described methods (Lisanby *et al*, 2003b; Moscrip *et al*, 2004; Spellman *et al*, 2009). For transport to the procedure room, subjects were sedated with i.m. ketamine (5.0–10 mg/kg) and xylazine (0.3–0.5 mg/kg) for the MST procedures or i.m. ketamine (3.0 mg/kg) and dexmedetomidine (0.075–0.15 mg/kg) for the ECT procedures. Xylazine was replaced with dexmedetomidine in the course of the study because of increasing tolerance of the subjects to xylazine. An additional dose of ketamine (2.5 mg/kg) was administered if needed to maintain sedation. Anesthesia and muscular paralysis were induced with i.v. methohexital (1.0 mg/kg) and succinylcholine (3.5 mg/kg), with a second bolus of each given at the initial dosages if the procedure could not be completed prior to anesthetic or paralytic emergence. Methohexital and succinylcholine are commonly used for clinical ECT (Bryson *et al*, 2012).

### Stimulation modalities

ECT was administered using a customized Spectrum 5000Q device (MECTA Corp., Tualatin, OR, USA) that allowed a wider range of stimulus current parameters. The electrical stimuli consisted of 0.2 ms bidirectional rectangular pulses delivered through electrodes 2.5 cm in diameter placed in the RUL position (Peterchev *et al*, 2015a) (Figure 1A). RUL electrode placement was used because it has fewer cognitive side effects than bilateral placement, and is often the first-line choice in clinical ECT (APA, 2001b; Sackeim *et al*, 2000; Sackeim *et al*, 2008).

MST was administered using a custom-made MagPro MST device (MagVenture A/S, Farum, Denmark). MST stimuli consisted of 0.36 ms cosine pulses delivered through a 10-cm-diameter coil placed on the vertex of the head (Peterchev *et al*, 2015a) (Figure 1B).

### Seizure threshold titration

ST was titrated using an ascending method of limits procedure (Peterchev *et al*, 2010b; Sackeim *et al*, 1987) utilizing successive stimulus trains with an increasing number of pulses, and hence train duration, while the other stimulus parameters (pulse amplitude, pulse width, and train frequency) were kept fixed.^2^ Amplitude was set to twice the individual amplitude-titrated ST determined at frequency of 50 pulses per second (pps) and 500 pulses in a prior study (Peterchev *et al*, 2015a) (range 230–565 mA for ECT and 66–86% of maximum stimulator output for MST; see Table 1 for individual amplitudes). Amplitude was selected in this manner in order to (1) compensate for individual anatomical variability and thus normalize the electric field exposure in the brain across subjects (Abbott *et al*, 2024; Deng *et al*, 2013; Peterchev *et al*, 2010b) and (2) provide electric field intensity in the brain that is comparable to that in clinical ECT as estimated by computational electric field models (Abbott *et al*, 2024; Lee *et al*, 2016, 2017).

ECT ST was titrated at 5, 10, 16, 25, 50, 100, and 240 pps. MST ST was titrated at 5, 10, 16, 25, and 50 pps for all subjects, and at 100 pps for two of the four subjects (MA and DY). Limitations of the MST device, namely its inability to stimulate at a high enough amplitude and frequency simultaneously, prevented titration at 100 pps in the other two subjects (CH and RZ) and at 240 pps for all subjects. MST data were acquired first, followed by the ECT data. ST data were collected from a total of 150 sessions.

We aimed to collect three ST estimates for each modality × frequency condition in separate sessions. Within modality, the order of the sessions with different frequency conditions was randomized. The first titration for each modality × frequency condition used coarse steps (70% increments in the number of pulses) to locate a range for the individual ST. Subsequent titrations were finer (30% pulse increments) to provide a more accurate estimate of the ST. Titration steps were separated by intervals of approximately 20 s. Seizure was determined by visual observation of tonic-clonic motor activity and concomitant electroencephalography (EEG) signal recorded by the MECTA Spectrum device. In instances where evidence of a seizure was ambiguous, i.e., if very brief (~ 1–2 s) motor or EEG seizure activity after the end of the stimulus was noted, the interval was lengthened to 60 seconds before the subsequent titration step. No more than six pulse trains were delivered in a single session. Sessions during which the subject had a seizure on the first titration step were discarded, as it is possible that the threshold was lower than what was observed. For 5 pps trains, since the seizure can take place completely during the stimulus train, strong tonic contractions during the stimulus train followed by motor suppression on the subsequent titration step were considered manifestation of a seizure. Sessions in which no clear seizure was identified were repeated.

### Seizure characteristics

The duration of the motor seizure was defined as the interval between the beginning of the stimulus and the end of the motor seizure expression. The one exception to this definition was for the 5 pps stimulation condition, since the motor expression typically started up to tens of seconds after the initiation of the stimulus, due to the slow rate of pulse delivery (see Supplementary Figure S6A). Therefore, for the 5 pps condition, we defined the beginning of the seizure when the motor expression started during the stimulus train. Ten sessions for which the seizure start was not noted were excluded from the seizure duration analysis.

The visually observed strength of the motor seizure expression in the four limbs (separate for tonic and clonic phase) and face was rated on a qualitative scale of “none-weak-medium-strong.” Electromyographic (EMG) recordings were also made for exploratory analyses (see Supplement).

### Electroencephalography

Brain activity during seizures was recorded with EEG using two bilateral fronto– mastoid channels through the MECTA Spectrum ECT device and software (gain = 5000, sampling rate = 140 Hz). Data processing was performed using MATLAB R2023 (MathWorks, Inc.). The EEG was bandpass filtered from 0.5–50 Hz, manually artifacted and resampled to 128 Hz. Pre-stimulus baseline activity was analyzed for 30 s before stimulus onset. Ictal activity was analyzed between the end of the stimulus train and the end of the EEG seizure or, if that was ambiguous, the end of the motor seizure. Artifact-free postictal activity was analyzed after the end of the seizure. Discrete wavelet transform was performed using the 4th-order Daubechies wavelet (Subasi, 2007). The EEG power was computed from the detail functions reconstructed from wavelet coefficients corresponding the following frequency bands: delta (δ: 0–4 Hz), theta (θ: 4–8 Hz), alpha (α: 8–16 Hz), and beta (β: 16–32 Hz). EEG power (in units of µV^2^/Hz) was log-transformed (base 10) and averaged over artifact-free 1-second epochs overlapping by 0.5 s.

EEG power analysis was carried out for 85 sessions; the rest were excluded due to malfunction of the recording computer (7), seizures shorter than stimulus duration (8, all at 5 pps), or insufficient duration of artifact-free seizure recording (50).

### Statistical analysis

The ST, seizure duration, and EEG power data were analyzed with linear mixed-effects models in JMP Pro 17 (SAS Institute Inc.) and plotted with JMP or MATLAB. Modality (ECT or MST), stimulus train frequency, and session were treated as fixed effects, and subject—as a random effect. Modality and subject were treated as nominal variables and session number as a continuous variable. Stimulus frequency was treated as a nominal variable to accommodate its strongly nonlinear effect on ST. The ST data, expressed as either number of pulses or train duration, were log-transformed to reduce heteroscedasticity across frequency conditions and improve normality. Statistical analysis of the EEG power also included EEG channel laterality (left, right), phase (baseline, ictal, postictal), and band (delta, theta, alpha, beta) as fixed effects. All fixed effects interactions were modeled; to allow this the 240 pps ECT condition was excluded since it does not have an MST counterpart. False discovery rate correction was used to account for multiple comparisons. The mixed-effects analysis was followed up with t-tests to compare ECT and MST as well as t-tests with Dunnett or Dunnett-Hsu multiple comparison correction to identify means significantly different from the minimum across stimulus frequency. Wald test was used to assess the significance of subject as a random variable. Results were considered significant for p < 0.05, unless otherwise noted.

## Results

### Seizure threshold

Seizure induction was generally reliable at all frequencies except for 5 pps. At 5 pps, seizures could not always be induced and the long duration and large number of the pulses led to premature termination of some titration sessions due to the subject’s emergence from anesthesia, MST coil heating, or ECT device errors.

Figure 2 shows ST versus stimulus frequency for ECT and MST. There was no significant interaction of modality × frequency (*F*_5,121_ = 0.808, *p* = 0.546), indicating that ST as a function of frequency behaved similarly for ECT and MST. The effect of session number was not significant either (*F*_2,121_ = 1.93, *p* = 0.200); this is expected since there was one session per week to minimize carryover effects, the frequency conditions were interleaved and randomized, and the seizure induction sessions were carried out regularly over the course of several years as part of a series of studies, so any lasting effects would have reached a steady state. Therefore, the interaction term and session number were removed from the final model. ST, quantified as the number of pulses required to induce a seizure, had a significant U-shaped dependence on stimulus frequency (Figure 2A; *F*_6,139_ = 68.9, *p* < .0001). Of the tested stimulus frequencies, the mixed-effects model identified 16 pps to minimize the number of pulses for seizure induction. This minimum ST did not differ significantly from the STs at 10 and 25 pps (*t*’s < 1.40, *p*’s > 0.564), but was significantly lower than the STs for 5, 50, 100, and 240 pps (*t*’s > 3.79, *p*’s < 0.0014). ST was affected by modality (*F*_1,139_ = 15.9, *p* = .0001), with MST having 18.8% lower ST than ECT, on average (*t* = −3.98, *p* = .0001). The least-squares estimates of the mean ST at 16 pps were 87.7 and 71.3 pulses for ECT and MST, respectively. There was no significant ST variance contributed by the individual subjects (*p* = 0.267), indicating that STs were consistent across subjects.

**Figure 2.**
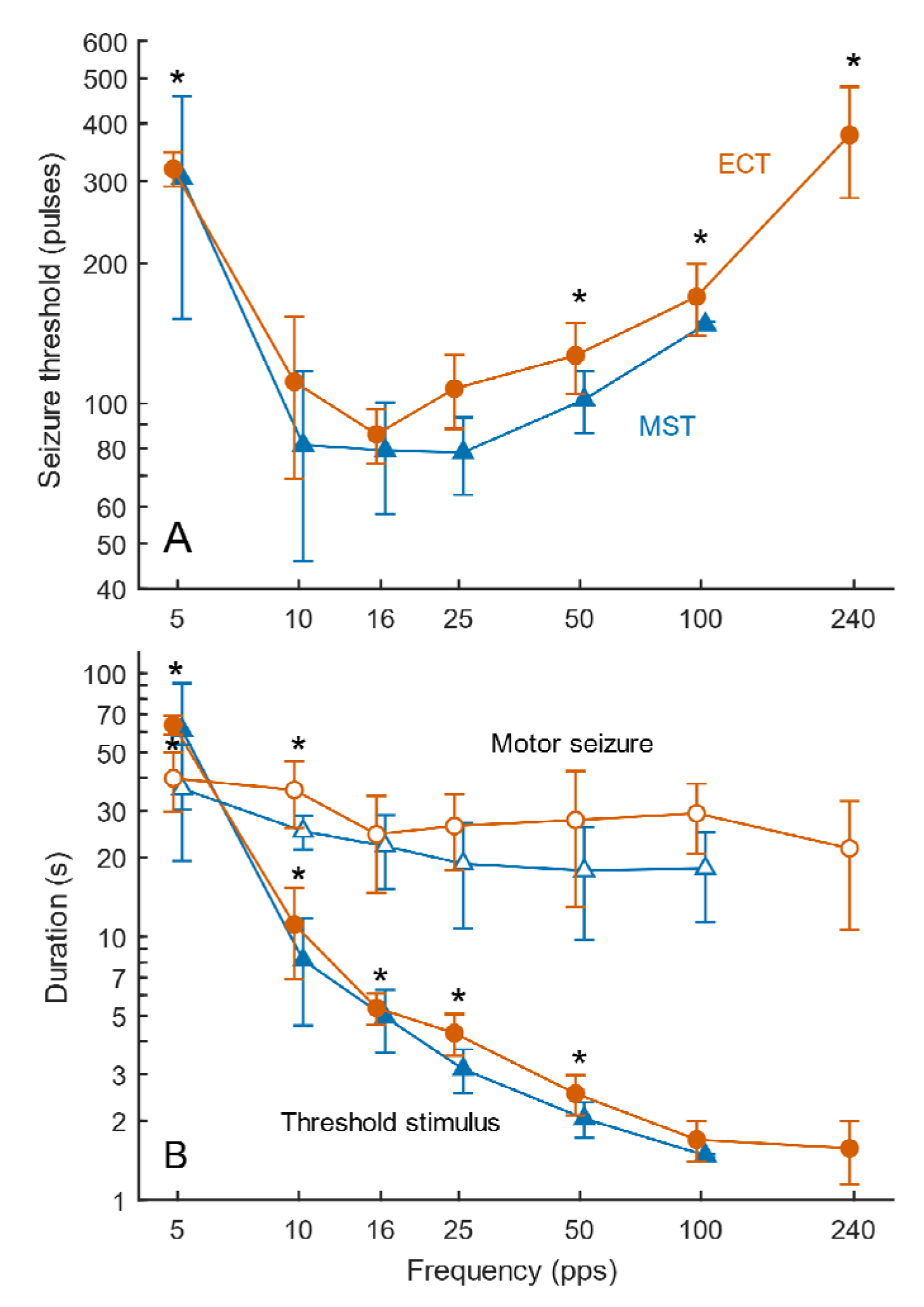
Seizure threshold (ST) and seizure duration for ECT (red) and MST (blue) as a function of stimulus pulse train frequency (pulses per second, pps). A: ST expressed as the number of pulses needed to induce a seizure at each frequency (stimulus total charge and energy are directly proportional to the number of pulses). B: Duration of the stimulus pulse train at ST (filled markers) and respective observed motor seizure duration (open markers). Markers and whiskers indicate the mean and standard deviation of the subject averages. The average for each subject was computed across all titration sessions at each frequency. Asterisks (*) mark levels significantly different (p < 0.01) from the respective minimum across frequencies. All axes are logarithmically spaced. Individual data points are shown in Supplementary Figures S1–S3.

Another way to quantify ST is by the duration of the stimulus train, which is equal to the number of pulses divided by the train frequency. As illustrated in Figure 2B, the train duration at ST decreased monotonically with increasing frequency (*F*_6,139_ = 378, *p* < .0001). The range of least-squares means spanned 63.4 s for 5 pps to 1.50 s for 240 pps for ECT. The mean train duration at 240 pps did not differ significantly from the one at 100 pps, but was significantly shorter than those for lower frequencies (*t*’s > 4.43, *p*’s < .0001).

### Seizure motor characteristics

Observed motor seizure expression strength in the unparalyzed arm was rated as “medium” for 88%–89% of the seizures (Supplementary Figures S4 and S5). Figure 2B shows the observed motor seizure duration for each stimulus frequency. There was no interaction between modality and stimulus frequency (*F*_5,111_ = 1.97, *p* = 0.118), indicating that stimulus frequency affected seizure duration similarly for ECT and MST. The effect of session number was not significant either (*F*_2,111_ = 1.76, *p* = 0.176). Therefore, the interaction term and session number were removed from the final model. Stimulus frequency significantly affected the motor seizure duration (*F*_6,129_ = 7.85, *p* < .0001). For ECT, the least-squares means of the motor seizure duration ranged from 42.1 s at 5 pps to 21.7 s at 240 pps. At 5 pps, the mean duration of the motor seizure was shorter than the duration of the respective stimulus trains, which exceeded 1 minute on average; motor seizure expression often started notably later than the train initiation and ended before or shortly after the end of the train. For higher stimulus frequencies (≥ 10 pps), seizure duration was longer than the stimulus train. Compared to the frequency with the shortest seizure duration (240 pps), only the 5 pps and 10 pps conditions had a significantly longer seizure duration (*t*’s > 3.76, p’s < .0013).

Modality had a significant effect on the motor seizure duration (*F*_1,129_ = 22.2, *p* < .0001). Motor seizure duration for MST was on average 26.9% shorter compared to ECT (*t* = −4.71, *p* < .0001).

Exploratory EMG recordings indicated that the ratio between tonic and clonic phase duration was comparable across stimulus frequencies (Supplementary Figure S8).

### Seizure EEG characteristics

EEG seizure duration was largely consistent with the observed motor seizure duration (Supplementary Figure S7). Baseline global power was significantly greater in the MST than in the ECT condition (*t* = 9.10, *p* < 0.0001; Supplementary Figure S9), likely reflecting differences in anesthetics used across these conditions. We subsequently normalized the ictal and postictal power relative to the baseline for each condition by subtracting the log-transformed baseline power from the log-transformed ictal and postictal power. Session number and EEG channel did not affect significantly the normalized global EEG power (*p*’s > 0.55). Consequently, we averaged the two channels within subject, condition, band, and session and did not model session number.

Figure 3 shows the average normalized EEG power data. There were significant main effects of seizure phase (*F*_1,130_ = 175, *p* < 0.0001) and stimulus frequency (*F*_5,133_ = 9.80, *p* < 0.0001). Further, significant interactions were frequency × phase (*F*_5,130_ = 8.34, *p* < 0.0001), frequency × modality (*F*_5,132_ = 4.96, *p* = 0.0006), and frequency × modality × phase (*F*_5,130_ = 4.17, *p* = 0.0021). Nonsignificant effects were modality (*F*_1,131_ = 1.67, *p* = 0.231) and modality × phase (*F*_1,130_ = 0.508, *p* = 0.477). There were also no significant differences between subjects (Wald test, p = 0.742). As expected, postictal power was lower than ictal power (*t* = −13.2, *p* < 0.0001). Comparing ictal power across stimulus frequencies, the 5 pps condition had significantly higher power than the minimum at 16 pps for ECT (*t* = 7.75, *p* < 0.0001), and the 5 pps and 10 pps conditions had a significantly higher power than the minimum at 25 pps for MST (*t* = 3.40, *p* = 0.0042 and *t* = 5.97, *p* < 0.0001, respectively). In the postictal phase, there were no significant differences in power across the stimulus frequencies relative to the condition with the least negative power (*p*’s > 0.112).

**Figure 3.**
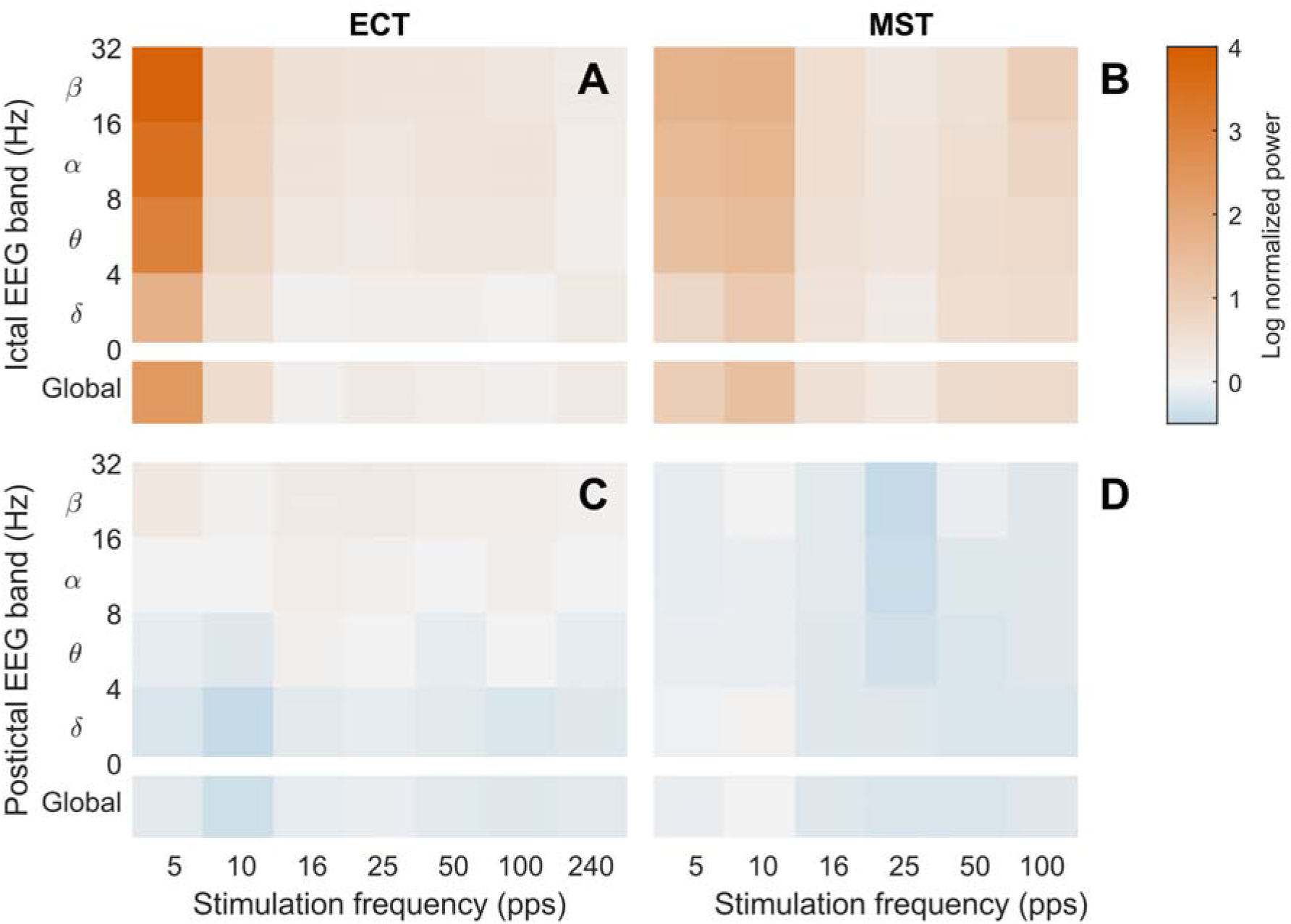
Average EEG spectral power of ictal (A, B) and postictal (C, D) period relative to baseline for ECT (left) and MST (right) for various stimulation frequencies. Bottom row within each subplot corresponds to global EEG power and rows above it correspond to discrete EEG bands (delta through beta). Individual raw and normalized data points are shown in Supplementary Figures S9–S12.

We further examined the effect of stimulation frequency on normalized EEG power within the delta, theta, alpha, and beta bands. In addition to the significant effects for global power, there were also significant main effects of band (*F*_3,533_ = 14.8, *p* < 0.0001) and modality (*F*_1,533_ = 8.83, *p* = 0.0031), and a significant modality × band interaction (*F*_3,533_ = 4.88, *p* = 0.0050). Notably, all interactions involving frequency × band were not significant (*p*’s > 0.591), indicating that the EEG power varied similarly with frequency across bands. Therefore, the significantly higher ictal power from low-frequency compared to high-frequency stimulation and the lack of significant frequency dependence in the postictal phase observed for the global power hold also across bands.

## Discussion

We showed that the number of pulses (and hence charge and energy) at ST varies as a function of the stimulus train frequency in a similar way for ECT and MST. The range of frequencies that results in the lowest ST is centered around 16 pps (equivalent to 8 Hz in the conventional ECT terms of pulse-pair frequency). This value is below the lowest frequencies, 10 Hz or 20 Hz, available in standard commercial ECT devices (MECTA Corp., 1997; SigmaStim, 2022; Somatics LLC, 2021). Our findings indicate that increasing stimulation frequency above 25 pps (50 Hz), as is typically done in conventional ECT titration, reduces seizure induction efficiency. This is consistent with several clinical studies that found a significant difference between ST at different stimulus train frequencies (Backhouse *et al*, 2018; Daskalakis *et al*, 2020; Girish *et al*, 2003; Swartz *et al*, 1989; Weaver *et al*, 1982).

Our analysis indicated that modality had a significant effect on ST. The higher ST for ECT than MST could be potentially explained by the fact that the ECT sessions used a modified sedation. Additionally, the individual pulse amplitudes for ECT were determined with a unidirectional pulse train (Peterchev *et al*, 2015a), whereas the trains used for frequency titration were bidirectional, and we have previously shown that bidirectional ECT trains can have higher ST than unidirectional trains (Spellman *et al*, 2009). Nonetheless, the lack of interaction between frequency and modality supports the understanding that the fundamental mechanism of seizure induction by ECT and MST is the same, specifically tetanic stimulation of neural populations with an induced electric field (Lee *et al*, 2016; Peterchev *et al*, 2012).

In our dosing paradigm, the stimulus amplitude was individually titrated (Peterchev *et al*, 2015a) and then doubled before ST was titrated by increasing the number of pulses. The absence of significant ST differences across subjects indicates that the pulse amplitude individualization compensated appropriately for individual differences and that additional titration of the number of pulses is unnecessary. Rather, a fixed number of pulses could be used for all subjects for a given modality and stimulus frequency. The clinical implications of this new approach to individualizing amplitude are just beginning to be explored (Abbott *et al*, 2024).

We observed reduction of the train duration at ST with increasing frequency, even though more pulses were delivered at higher frequencies (Figure 2B). Similar observations in other nonhuman animal studies led some researchers to conclude that stimulus frequencies above 100 pps induce seizures more efficiently than lower frequencies (Hovorka *et al*, 1960; Hyrman *et al*, 1985; Liberson, 1945; Weaver *et al*, 1974). These results contradict human ECT and MST studies that suggest lower frequencies are more efficient (Backhouse *et al*, 2018; Daskalakis *et al*, 2020; Devanand *et al*, 1998; Girish *et al*, 2003; Swartz *et al*, 1989; Weaver *et al*, 1982). This discrepancy may stem from the fact that in some of the animal studies frequency and number of pulses were varied simultaneously since frequency was evaluated for a fixed train duration (Hovorka *et al*, 1960; Liberson, 1945; Woodbury and Swinyard, 1952). Thus, an increase in frequency was always associated with a proportional increase of the number of pulses. The probability of a seizure increases as more pulses are delivered (Alexander, 1953; Peterchev *et al*, 2010b; Peterchev *et al*, 2016); consequently, higher frequencies will be biased to lower ST. Nevertheless, these prior studies contain clues that lower frequencies may be more efficient if the number of pulses were held constant, as the simultaneous increase of both frequency and number of pulses produced slowing incremental reduction of ST, and eventually the threshold increased at high frequencies (>> 300 pps).

The EEG data indicate that while the seizure strength was overall comparable between ECT and MST, stimulus frequency significantly affected seizure expression and did so in a different way for ECT and MST. The key finding is that the lowest stimulus frequencies (5 pps and 10 pps) produced seizures with higher ictal power than higher stimulus frequencies. One limitation of this observation is that at 5 pps the duration of the stimulus train typically exceeded the duration of the seizure (see Figure 2B), and therefore the seizures that were recorded were markedly longer and perhaps more robust than the average. Nonetheless, this was not the case for the 10 pps condition which produced the highest ictal power for MST, and moreover the individual seizures with highest power were recorded for the 5 pps and 10 pps conditions. Also, the differences in seizure expression between ECT and MST may be confounded by the anesthesia differences, reflected in the baseline power, although prior studies of nonhuman primates have reported differences in seizure EEG expression under the same anesthesia as well (see Supplementary Figure S13) (Cycowicz *et al*, 2018). Other factors that distinguish ECT and MST and could contribute to seizure expression differences are the induced electric field distribution and the pulse shape and duration (Deng *et al*, 2011). For example, compared to the symmetric bilateral electric field of circular-coil-on-vertex MST, RUL ECT stimulates stronger in the right hemisphere and in depth (Lee *et al*, 2017).

The present study suggests stimulus trains in the 10−25 pps range are more effective at generating an excitatory state that overcomes innate cerebral inhibition than trains at higher or lower frequencies. While we did not probe the underlying mechanisms, this optimal stimulation frequency range can induce prolonged depolarization and epileptiform activity in hippocampal slices (Bracci *et al*, 2001; Thompson and Gahwiler, 1989) and coincides with a 10 Hz “recruiting rhythm” observed with EEG during endogenous seizure onset (Gastaut and Broughton, 1972). In contrast, higher stimulus frequencies can suppress ongoing seizure activity by extending the neural refractory period, inducing intermittent axonal block, and desynchronizing firing patterns (Feng *et al*, 2014; Wilson and Moehlis, 2015). Lower frequencies (~ 1 pps) can suppress seizures as well by inducing long-lasting hyperpolarization (Toprani and Durand, 2013), suggesting possible explanations of the U-shaped curve we observed in Figure 2A. Our EEG analysis also found frequency affected ictal expression, suggesting shifts in the dynamics of neural inhibitory and excitatory processes with stimulation frequency. An extended discussion of putative mechanisms is provided in the Supplement.

Our results, coupled with prior reports, suggest that when individualizing the ECT or MST dose it is most efficient to maximize the stimulus train duration first, and only then increase the train frequency. This dosing/titration approach may be especially advantageous for patients with a high ST, which may otherwise exceed the FDA-regulated device limit of 100 J (MECTA Corp., 1997; SigmaStim, 2022; Somatics LLC, 2021).

Since this was a study in nonhuman primates focused on seizure induction efficiency, we cannot make conclusions regarding clinical implications for efficacy or side effects. Few studies have investigated the impact of frequency on clinical outcomes (Daskalakis *et al*, 2020; Kotresh *et al*, 2004; Roepke *et al*, 2011; Weissman *et al*, 2020), and some of these have confounds (e.g., mismatch in the number of pulses across frequency conditions). Trials will be needed to determine if 10–25 pps results in more efficient seizure induction in patients, and to examine the impact on clinical outcomes.

## Data Availability

Individual data used in the table, figures, and statistical analyses are available from the Duke Research Data Repository (doi: https://doi.org/10.7924/r4765pt3p)

## Acknowledgments

The authors thank Mohamed Aly, Niko Reyes, and Brian Chan for assisting in the experimental procedures and Won Hee Lee for generating Figure 1. The data presented here were published in part and with preliminary analyses as conference abstracts at the 19^th^ and 31^st^ Annual Meeting of the International Society for ECT and Neurostimulation (Feuer *et al*, 2022; Peterchev *et al*, 2010a) and Society of Biological Psychiatry 70^th^ and 76^th^ Annual Scientific Convention (Peterchev *et al*, 2015b).

## Author contributions

Angel V. Peterchev: conceptualization, methodology, formal analysis, investigation, resources, data curation, writing - original draft, writing - review & editing, visualization, supervision, project administration, funding acquisition; Zhi-De Deng: formal analysis, data curation, writing – review & editing. Christopher Sikes-Keilp: investigation, data curation, writing - original draft; Elyssa Feuer: formal analysis, data curation, writing – review & editing. Moacyr A. Rosa: investigation; and Sarah H. Lisanby: conceptualization, methodology, resources, writing - original draft, writing - review & editing, supervision, project administration, funding acquisition.

## Footnotes

1 In ECT the stimulus frequency is conventionally defined as the repetition rate of pairs of pulses of opposite polarity measured in hertz (Hz). In contrast, the MST stimulus frequency is defined as the repetition rate of individual pulses, again in Hz. For consistency, in this paper we refer to the repetition rate of individual pulses reported as pulses per second (pps), unless otherwise noted.

2 The total stimulus charge or energy—common dose metrics in ECT—are directly proportional to the number of pulses.

