## Supplementary material for "Optimal Frequency for Seizure Induction with Electroconvulsive Therapy and Magnetic Seizure Therapy"

### Supplement

Angel V. Peterchev, PhD[1,2,3,4,*], Zhi-De Deng, PhD[5], Christopher Sikes-Keilp, MD[1,6], Elyssa C. Feuer, BA[7], Moacyr A. Rosa, MD PhD[1,8], and Sarah H. Lisanby, MD[5]

[1]Department of Psychiatry and Behavioral Sciences, Duke University, Durham, NC, USA

[2]Department of Biomedical Engineering, Duke University, Durham, NC, USA

[3]Department of Electrical and Computer Engineering, Duke University, Durham, NC, USA

[4]Department of Neurosurgery, Duke University, Durham, NC, USA

[5]Noninvasive Neuromodulation Unit, Experimental Therapeutics and Pathophysiology Branch, Intramural Research Program, National Institute of Mental Health, National Institutes of Health, Bethesda, MD, USA

[6]Department of Psychiatry, University of North Carolina, Chapel Hill, NC, USA

[7]University of Toledo College of Medicine and Life Sciences, Toledo, OH, USA

[8]Institute for Advanced Research in Neurostimulation, São Paulo, SP, Brazil

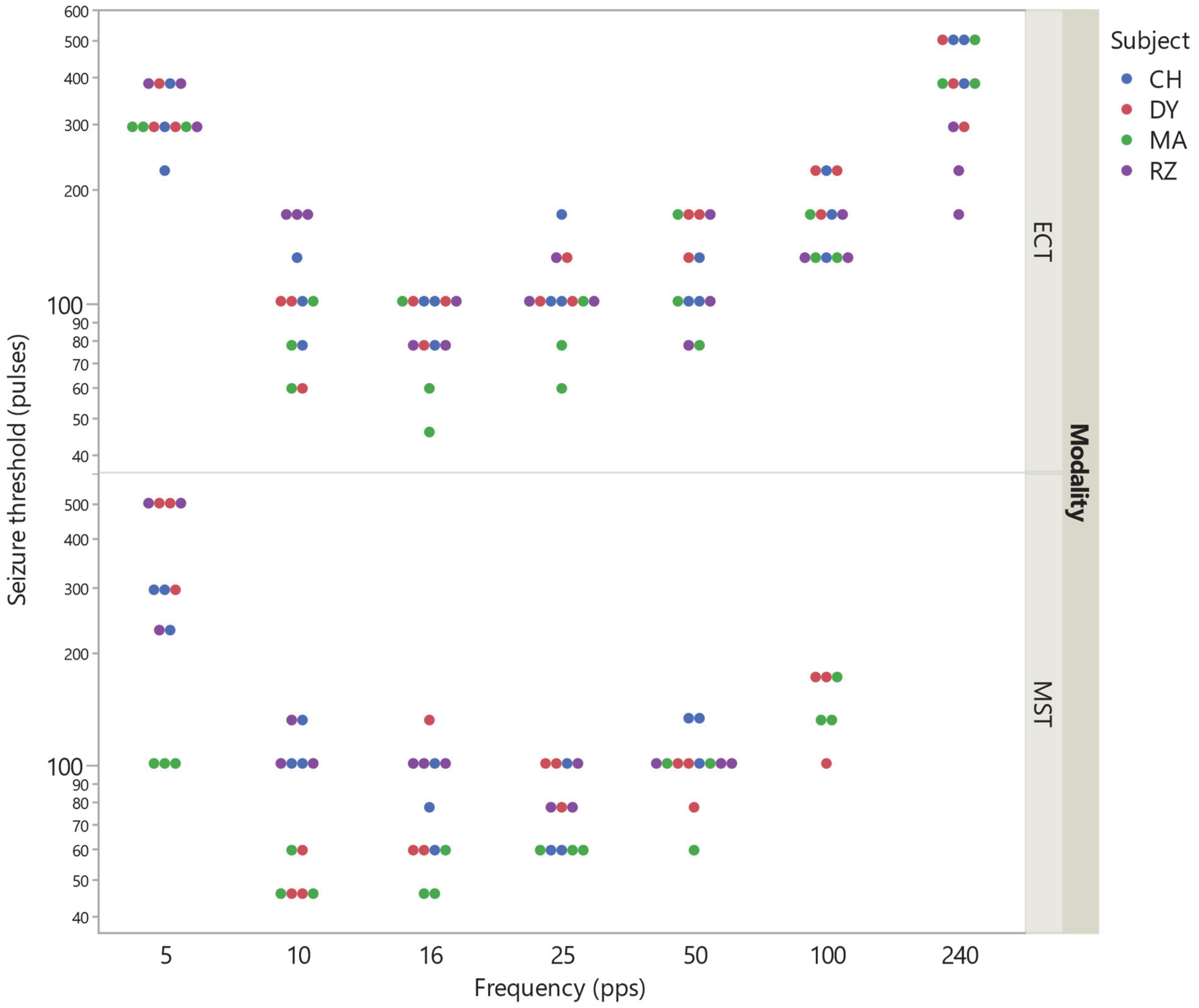


**Figure S1.** Individual seizure threshold (number of pulses) data corresponding to Figure 2A.

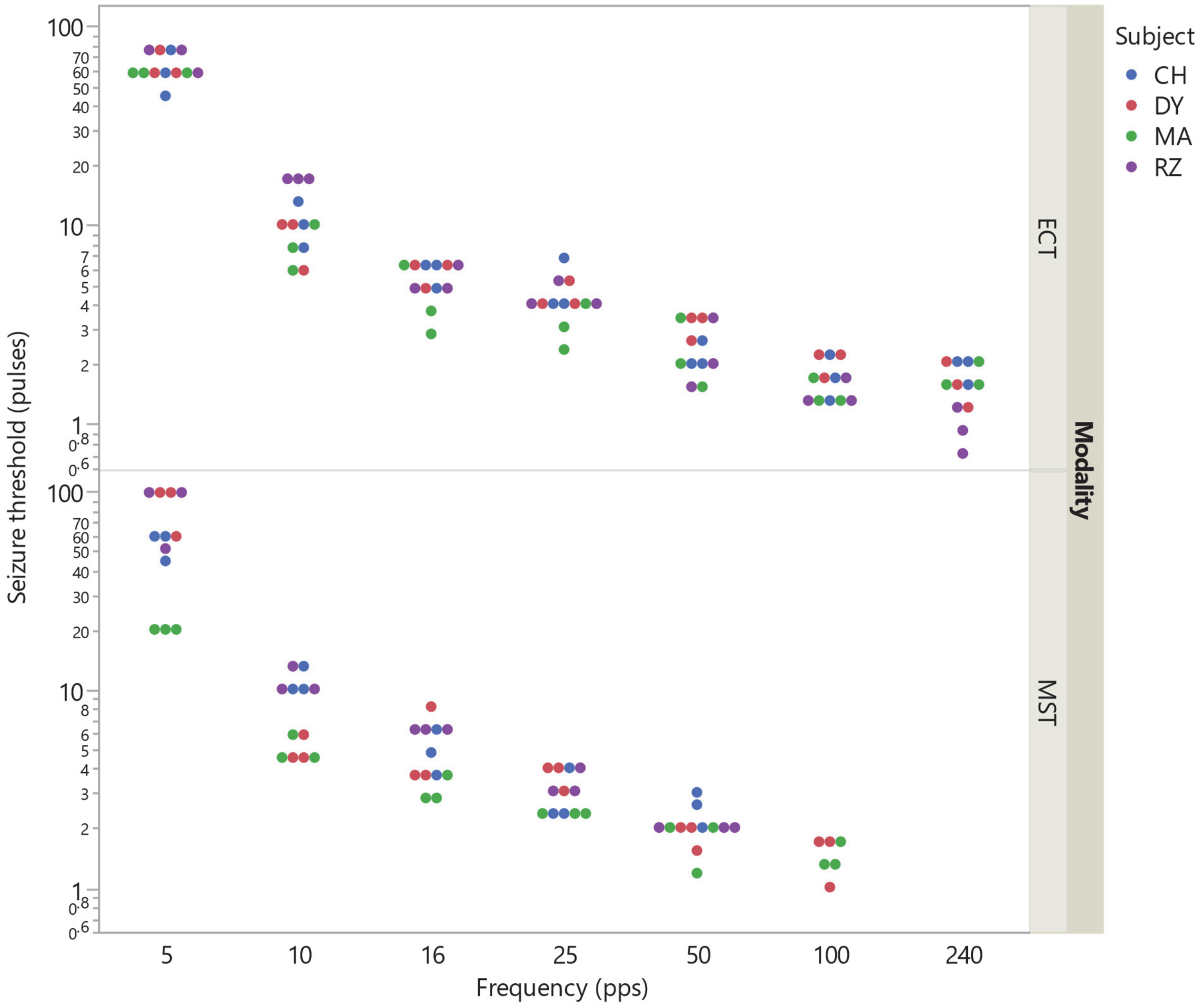


**Figure S2.** Individual seizure threshold (train duration) data corresponding to Figure 2B.

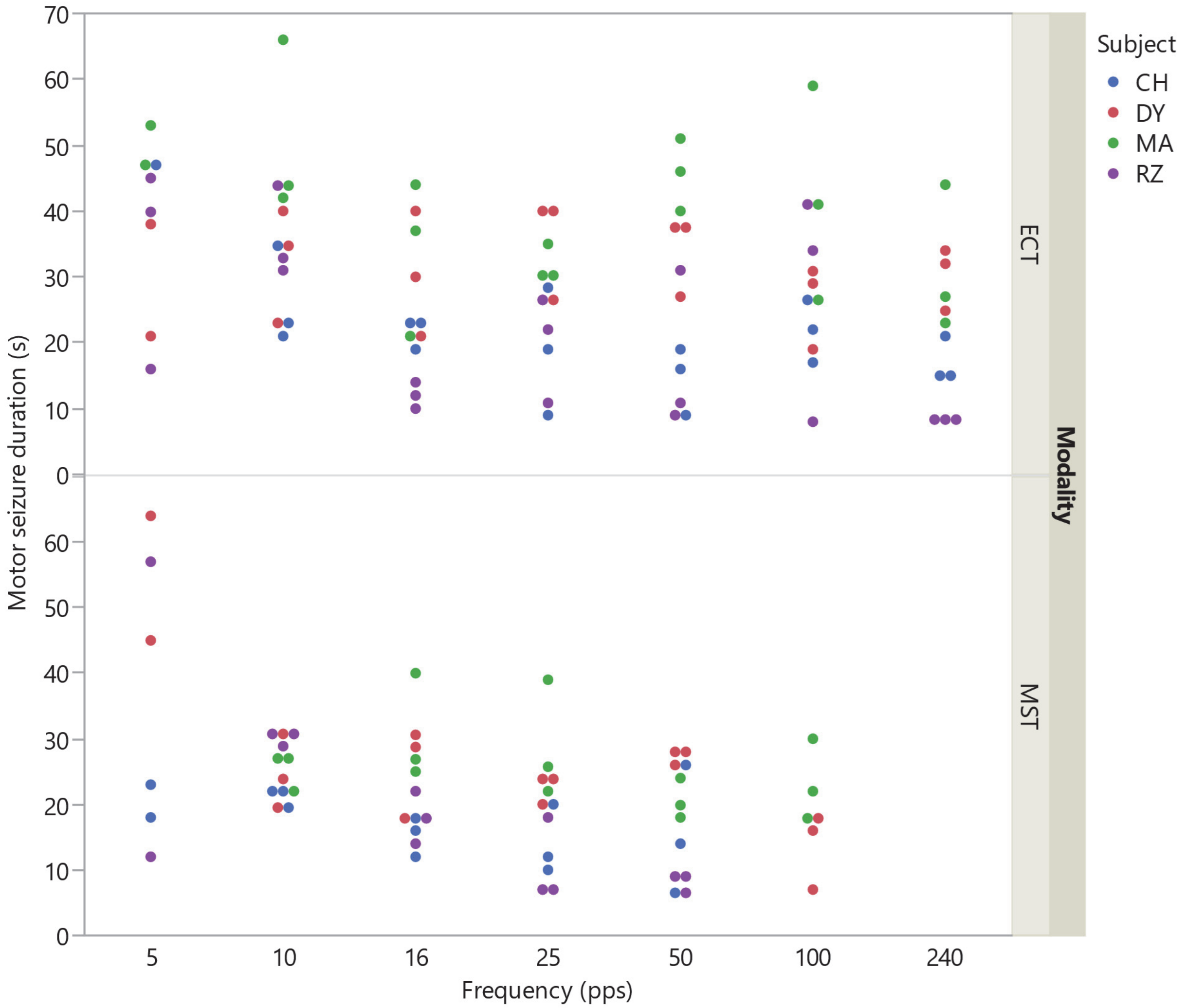


**Figure S3.** Individual motor seizure duration data corresponding to Figure 2B.

### Qualitative seizure expression rating

*Seizure rating methods*

The visually observed strength of the motor seizure expression in the four limbs (separate for tonic and clonic phase) and face was rated on a qualitative scale of “none-weak-medium-strong.” Seizure strength ratings were documented for 143 of the 150 sessions. These qualitative ratings are reported only descriptively, since they were not standardized and were conducted by various raters.

*Seizure rating results*

As expected, in the paralyzed limbs observed seizure strength was rated as “none” in the vast majority of sessions (94%–96%). In the face, 25.5%, 52.5%, and 22.0% of the seizures were rated as “none”, “weak”, and “medium”, respectively. These ratings likely depend strongly on the depth of paralysis. In contrast, observed seizure strength in the unparalyzed arm was rated as “medium” for the vast majority (88%–89%) of seizures, as shown in Figures S4 and S5.

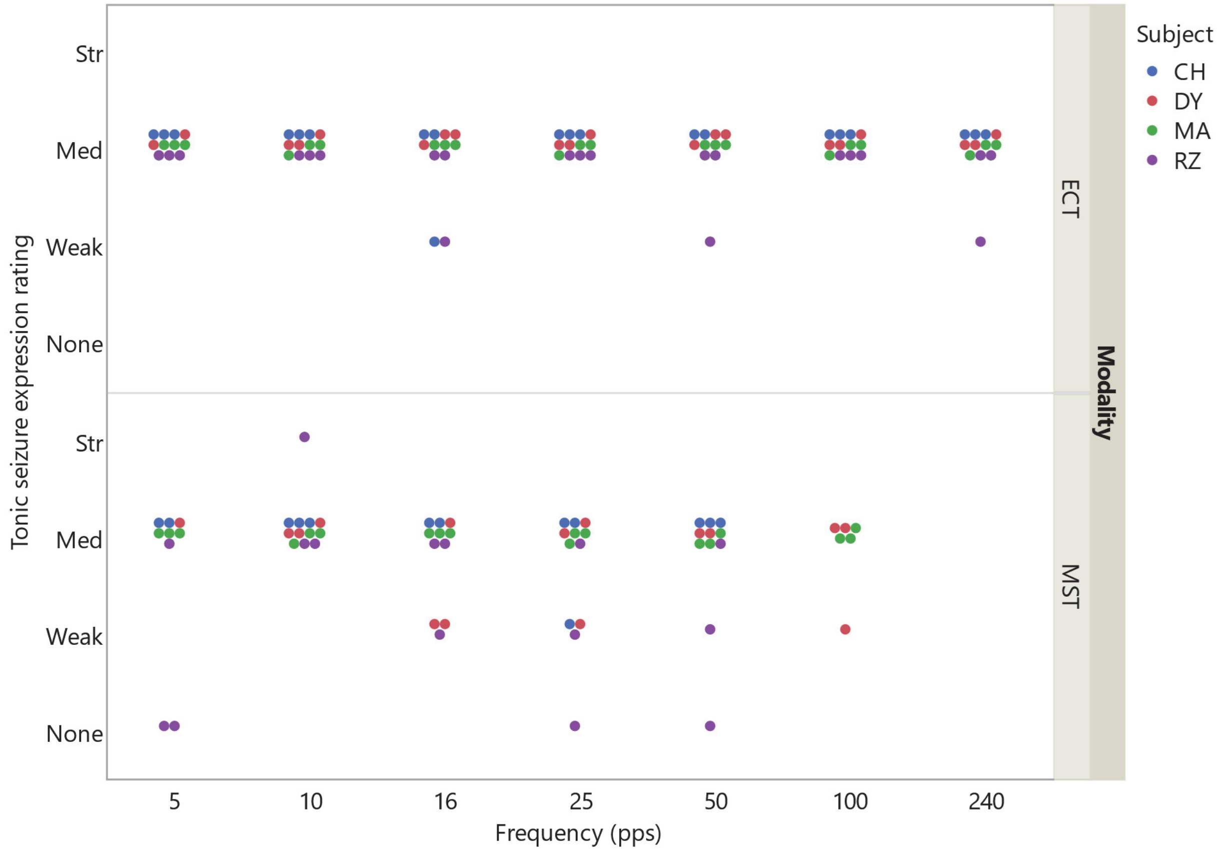


**Figure S4.** Tonic motor seizure expression rating for unparalyzed arm.

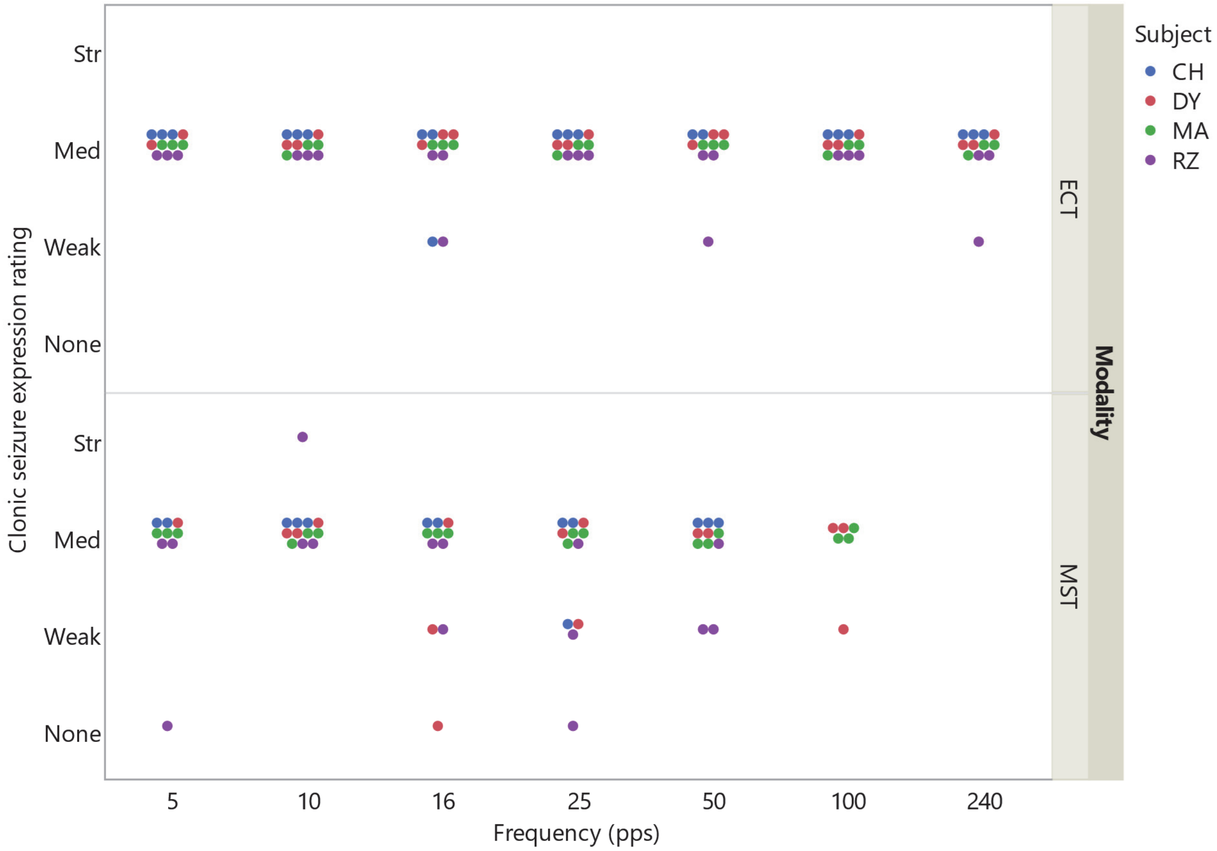


**Figure S5.** Clonic motor seizure expression rating for unparalyzed arm.

### Electromyography (EMG)

*EMG methods*

Unlike EEG data, EMG signals can be collected and analyzed during the ECT and MST stimulus delivery, providing additional quantitative information about the seizure characteristics. EMG data collection was piloted during 17 of the ECT sessions from all seven stimulus frequency conditions (5 pps, n = 2; 10 pps, n = 2; 16 pps, n = 4; 25 pps, n = 2; 50 pps, n = 3; 100 pps, n = 2; 240 pps, n = 2). EMG signal was recorded with needle electrodes inserted in the first dorsal interosseous muscle, amplified with bioamp (BIOAMP-4, SA Instrumentation Co.), and digitized at 5 kHz sampling frequency (USB-6009, National Instruments). We extracted the EMG signal from 5 seconds before stimulus onset to 5 seconds after seizure termination. Tonic and clonic seizure duration was calculated based on methods of Conradsen et al. (Conradsen *et al*, 2013). Briefly, the EMG was resampled to 1024 Hz and decomposed into a high-frequency (HF) component (64–256 Hz) and a low-frequency (LF) component (2–8 Hz) using an 8-level decomposition with the Daubechies 20 (db20) wavelet. The HF/LF ratio was computed. The tonic-clonic transition was defined as the time where the HF/LF ratio drops to below 20% of its maximum (Figure S6).

*EMG results*

Seizure duration determined by EMG was consistent with the observation of motor activity (Figure S7A). There was a significant effect of stimulus frequency on tonic phase duration (log-transformed frequency, $R^2 = 0.297$, $p = 0.0238$), but the tonic phase proportion of the total seizure length did not change significantly with stimulus frequencies (Figure S8). This suggests that although frequency did affect the overall seizure duration, the split of the seizure in tonic and clonic components was not altered significantly.

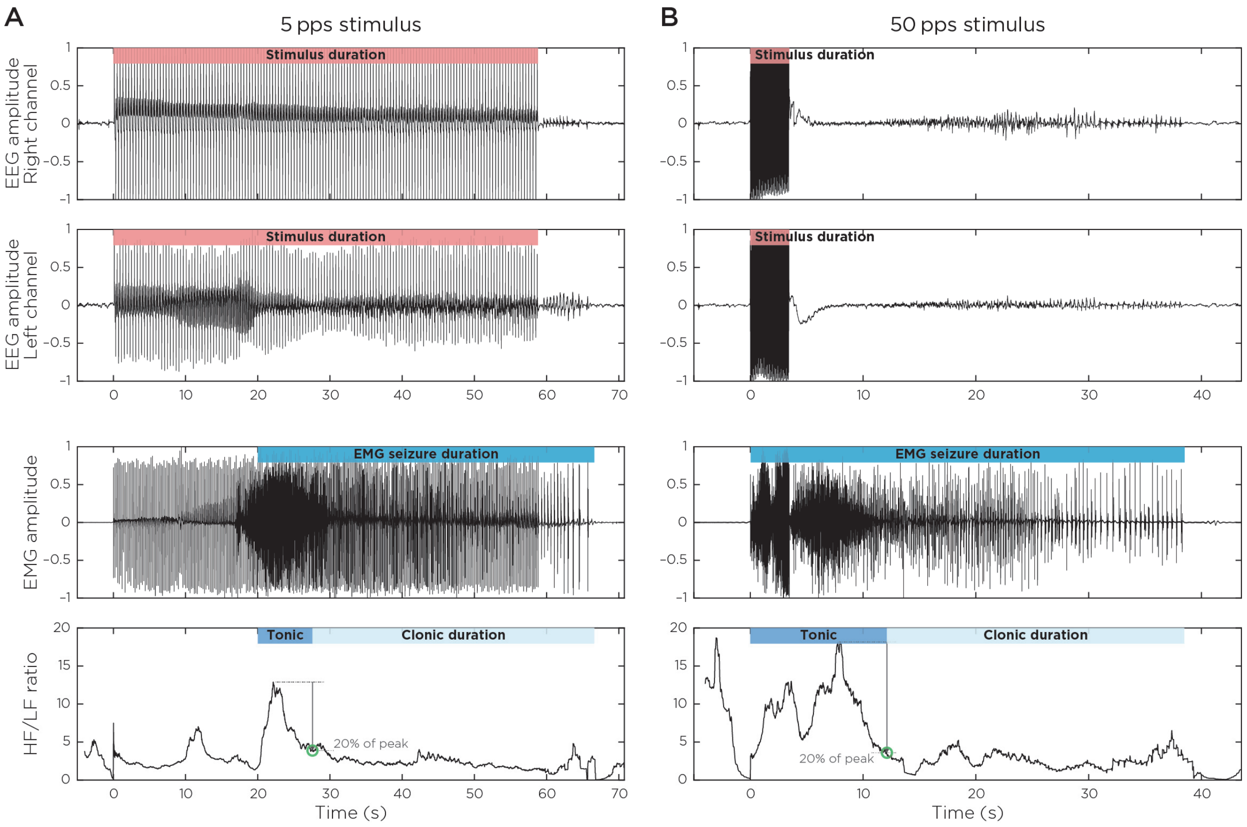


**Figure S6.** EMG recorded from one subject (MA) receiving ECT with (A) 5 pps and (B) 50 pps stimulus frequency. The top two rows show right and left normalized EEG channel amplitude. The third row shows normalized EMG signal from 5 seconds before stimulus onset to 5 seconds after seizure termination. The bottom panels show the corresponding high-frequency-to-low-frequency (HF/LF) EMG ratios. In the bottom panel, the blue bar differentiates the tonic and clonic seizure phases.

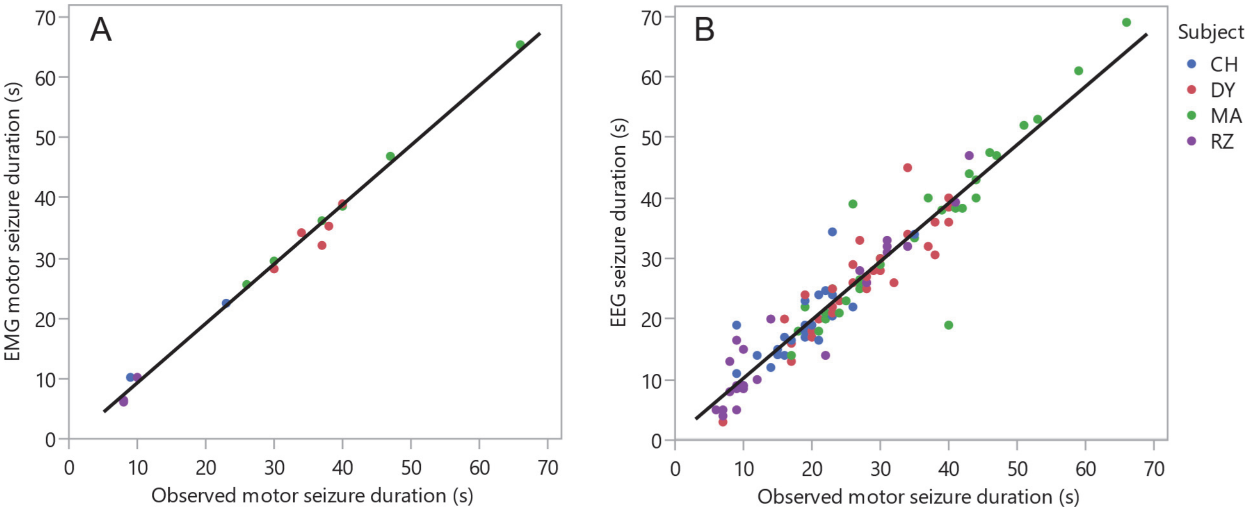


**Figure S7.** Correlation between observed motor seizure duration and (A) EMG motor seizure duration ($R^2 = 0.992$, $p < 0.0001$) and (B) EEG seizure duration ($R^2 = 0.899$, $p < 0.0001$). Colors denote subject and regression line is in black.

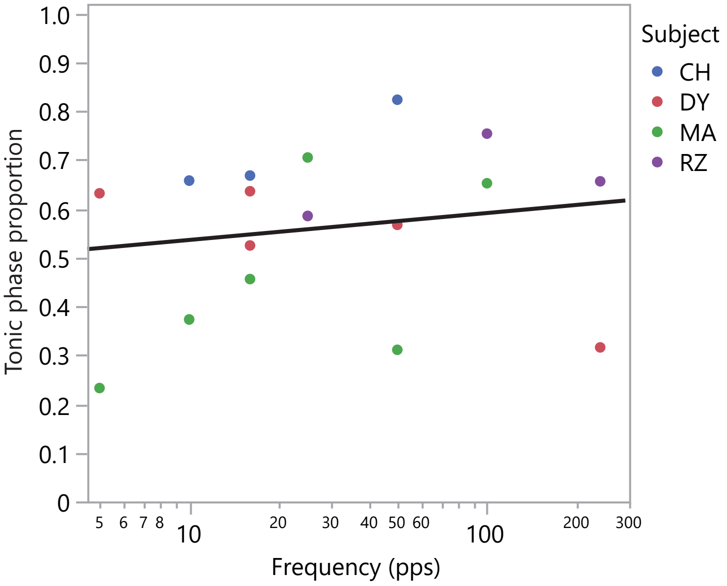


**Figure S8.** The tonic phase proportion relative to total EMG seizure duration did not vary significantly across stimulus frequency ($R^2 = 0.0288$, $p = 0.515$). Colors denote subject and regression line is in black.

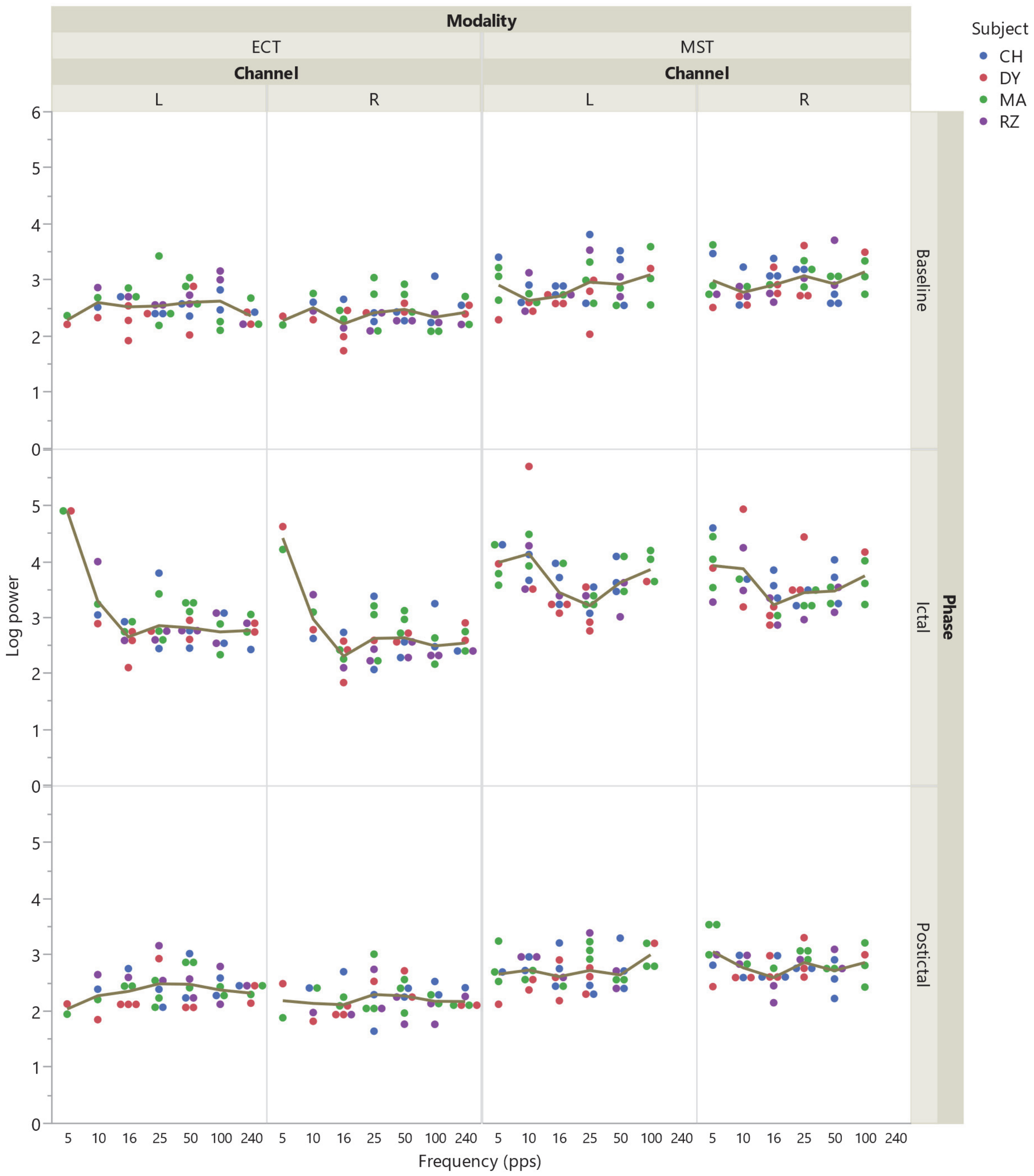


**Figure S9.** Log-transformed global EEG power across stimulation conditions (modality and frequency), EEG channels (left, L, and right, R), and seizure phase (baseline, ictal, and postictal). Colored dots correspond to data from individual sessions and subjects. Lines show the average within stimulation frequency.

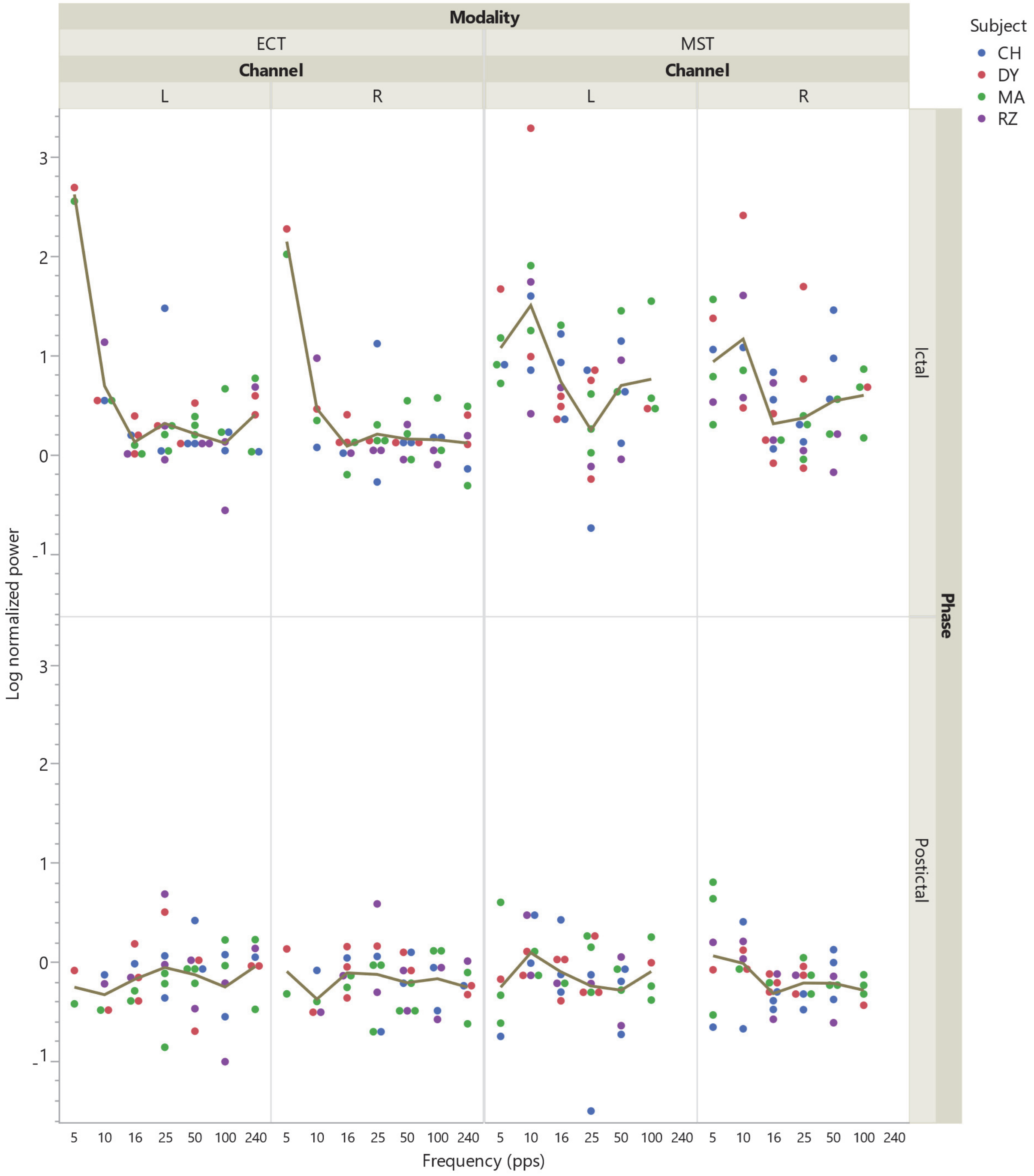


**Figure S10.** Log-transformed global EEG power normalized to baseline across stimulation conditions, EEG channels, and seizure phase. Display conventions as in Figure S9.

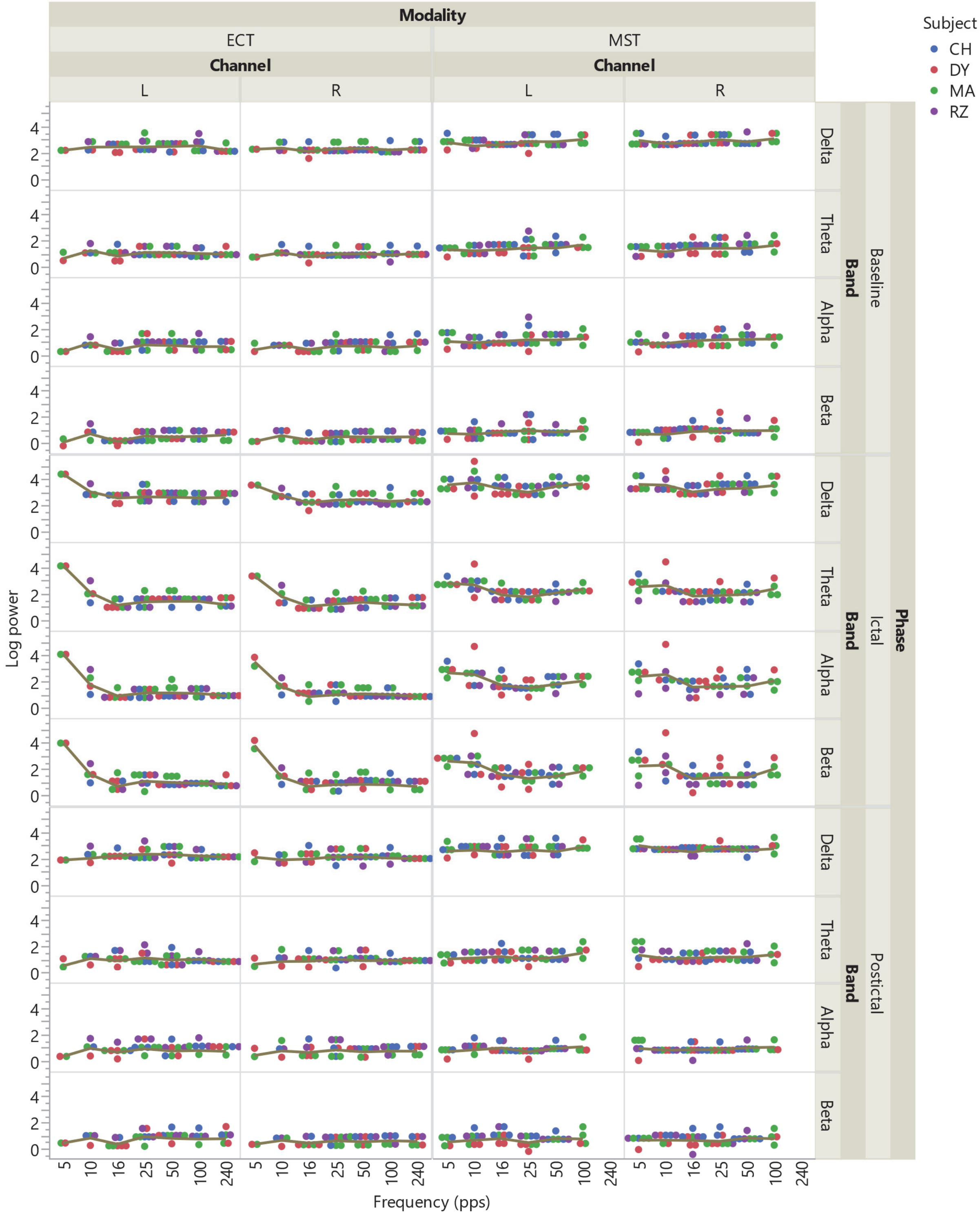


**Figure S11.** Log-transformed EEG power across stimulation conditions, EEG channels, seizure phase, and EEG bands. Display conventions as in Figure S9.

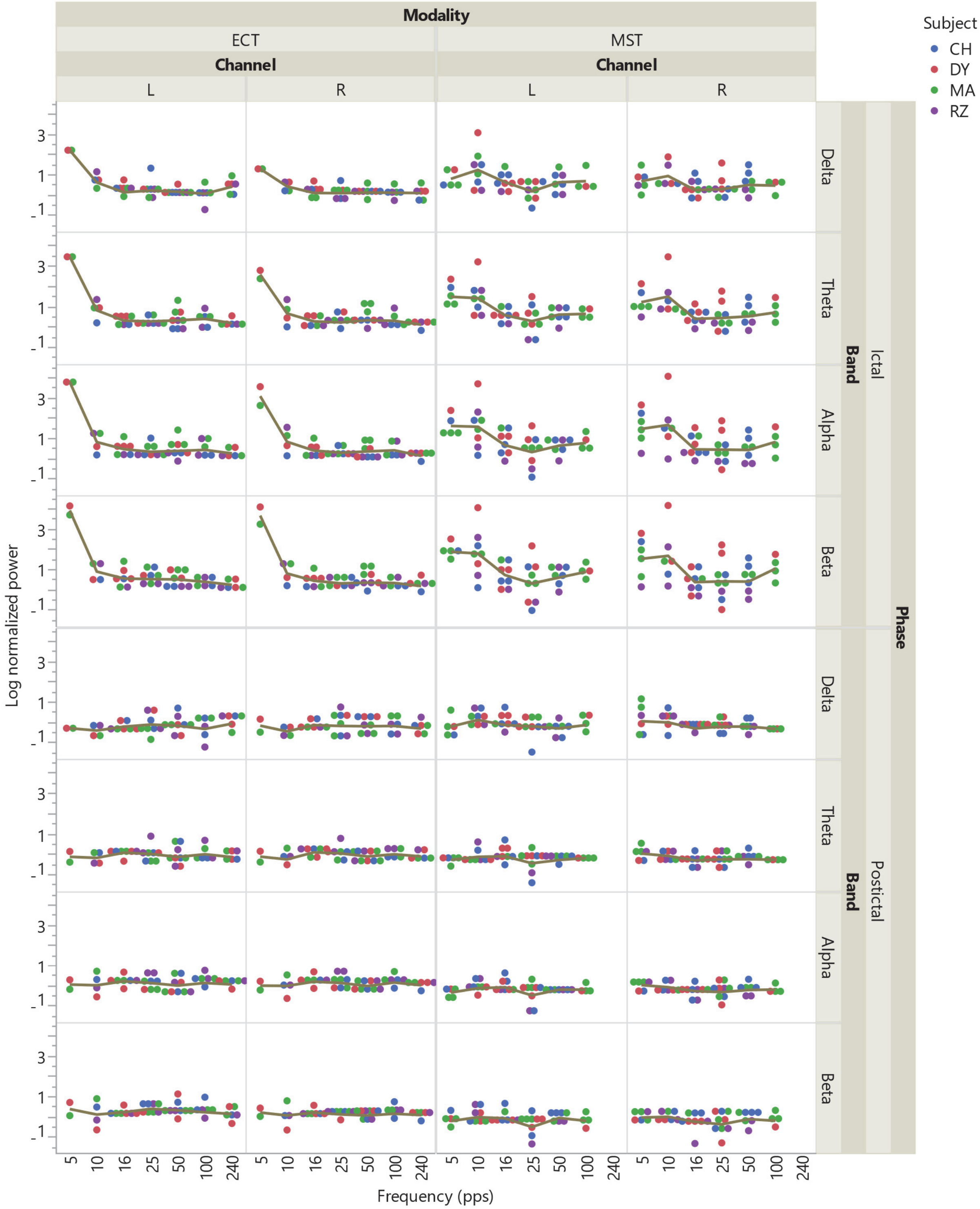


**Figure S12.** Log-transformed EEG power normalized to baseline across stimulation conditions, EEG channels, seizure phase, and EEG bands. Display conventions as in Figure S9.

### Additional discussion

*EEG comparison to prior study*

Differences in seizure expression between ECT and MST in nonhuman primates have been discussed previously. Cycowicz et al. (2018) reported a randomized experiment where 24 macaca mulatta (12 male) received 6 weeks of daily 50 Hz (100 pps) ECS or 50 Hz (50 pps) MST (Cycowicz *et al*, 2018). Threshold seizure induction resulted in differential EEG power distribution during the ictal and postictal periods. Specifically, ictal EEG power for ECT increased progressively from the low (delta) to high (beta) frequency bands; MST induced seizures had higher delta power, but lower beta power compared to ECS seizures. In the postictal period, ECT produced robust suppression of EEG power in the delta-to-alpha frequency bands, but less suppression of beta power, while MST produced slight suppression of theta-to-beta power, but no suppression of delta power (Cycowicz *et al*, 2018). Figure S13 compares our data with those of Cycowicz et al. In the present work, ECT was delivered at individualized amplitude of 364 ± 148 mA, less than half the amplitude (800 mA) used by Cycowicz et al. Further differences between the present study and Cycowicz et al. included the electrode placement (unilateral versus bilateral), shorter pulse width (0.2 ms versus 0.5 ms), and older nonhuman primate subjects (13.32 ± 2.73 years versus 2.83 ± 0.46 years). Compared to Cycowicz et al., MST was also delivered at a different amplitude in the present study (78 ± 9% of maximum device output, MSO, compared to 100% MSO) and with a different device (MagPro versus Magstim). Presumably due to these differences, the ECT ictal power and postictal suppression are greatly reduced in the present work, whereas the MST ictal seizure expression is reduced to a lesser degree and the postictal suppression is increased. Nonetheless, the different shape of the ictal power distribution across the EEG bands between ECT and MST is similar between the two studies, suggesting potential mechanistic differences between the two modalities.

Finally, both datasets in Figure S13 are at seizure threshold (ST), whereas clinical treatments are conventionally administered above ST. The present study did not explore suprathreshold stimulation, while Cycowicz et al. did, finding that increasing the train duration, and hence the number of pulses, by 150% relative to the ST, enhanced the ictal EEG power in the theta, alpha, and beta band and post-ictal suppression in the theta and alpha band for ECT but not MST. This points to potentially important further interactions between stimulation modality and the stimulus parameters.

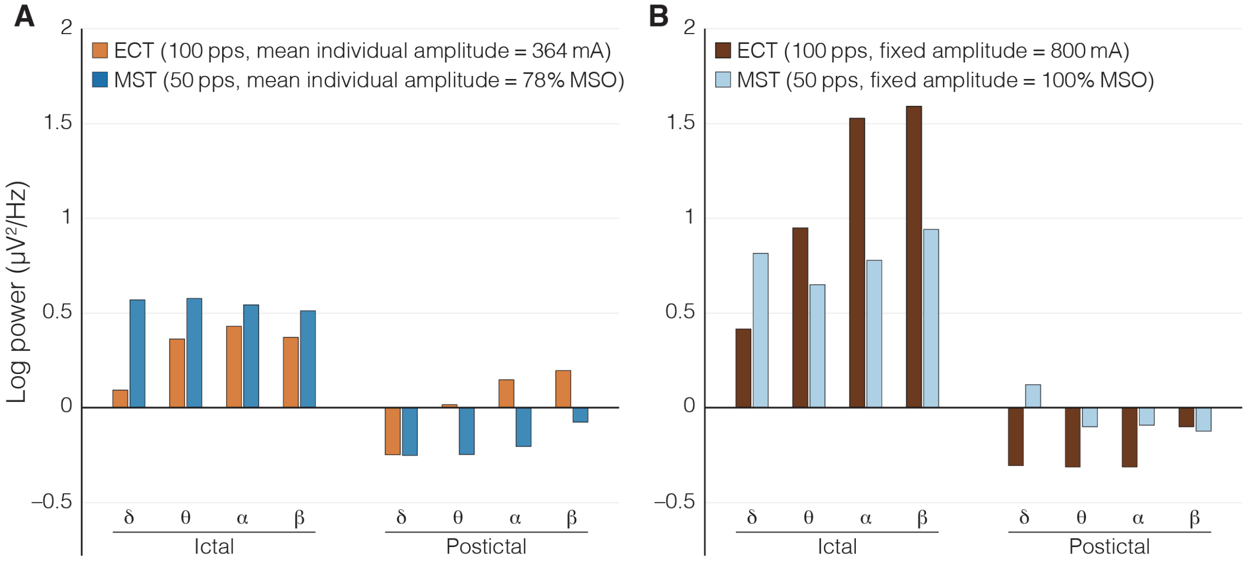


**Figure S13.** Ictal and postictal EEG power normalized to pre-stimulus baseline power for ECT with 100 pps and MST with 50 pps: A) The present work and B) Cycowicz et al., 2018 (Cycowicz *et al*, 2018). In the present work, ECT was delivered at individualized amplitude of 364 ± 148 mA; MST was delivered at individualized amplitude of 78 ± 9% maximum stimulator output (MSO). Cycowicz et al. delivered ECT at fixed amplitude of 800 mA and MST at fixed amplitude of 100% MSO.

*Putative physiological mechanisms*

We demonstrated that stimulus trains in the 10−25 pps range are most efficient for inducing seizures, and the ST increases sharply at lower frequencies and gradually at higher frequencies. This finding is largely consistent with evidence from in vitro, animal, clinical, and computational studies that offer insights into potential mechanisms. Electrical stimulation of rat hippocampal slices with 10 pps for 2–3 s preceded by 40 pps for 1 s (Bracci *et al*, 2001) or 100 pps for 0.2–0.4 s (Bracci *et al*, 1999; Kaila *et al*, 1997) results in prolonged depolarization of pyramidal cells. Longer stimulation with 3–10 pps for 30–60 s (Thompson and Gahwiler, 1989) or 60 pps for 2 s (Stasheff *et al*, 1985) can trigger epileptiform activity. The proposed mechanism was a reduction in GABA-A-mediated inhibitory postsynaptic potentials associated with an increase in the concentration of extracellular K+ and intracellular Cl− (Bracci *et al*, 1999, 2001;

Kaila *et al*, 1997; Thompson *et al*, 1989). Increased extracellular K+ concentration attenuates hyperpolarizing/inhibitory currents in surrounding neurons, and increased intracellular Cl− concentration causes a switch of GABA-A receptor function from inhibitory to excitatory (American Epilepsy Society, 2006; Boison, 2013). Further, changes in K+ can trigger transitions between different neural activity patterns, including from resting state to single spike firing or to depolarization block (Barreto and Cressman, 2011; Contreras *et al*, 2021; Wu and Shuai, 2012). Very low stimulus frequencies (< 1 Hz) produce only small neuromodulatory effects in the brain (Julkunen *et al*, 2012; Pellicciari *et al*, 2016), and there is an opportunity for inhibitory currents to operate to restore equilibrium (Toprani and Durand, 2013).

Another important observation is that electrical stimulation at both low (~ 1 pps) or high frequencies (~ 130 pps) can suppress ongoing epileptic activity (Durand and Bikson, 2001; Toprani *et al*, 2013; Yu *et al*, 2018). Repetitive transcranial magnetic stimulation (rTMS) pulse trains delivered at 1 pps are inhibitory (Di Lazzaro *et al*, 2011), and rTMS at 1 or 5 pps attenuated several EEG markers of penicillin-induced seizure activity in rats, while trains delivered at 10 pps facilitated the seizure (Lin *et al*, 2014). The anti-seizure mechanisms of low frequency stimulation involve long-lasting hyperpolarization mediated by GABA-B inhibitory postsynaptic potentials and slow afterhyperpolarization (Toprani *et al*, 2013). Putative mechanisms of high frequency stimulation are extensions of the refractory period, normally less than 2 ms in central neurons (Jankowska *et al*, 2022), and induction of intermittent axonal block and neural firing desynchronization (Feng *et al*, 2014; Guo *et al*, 2018; Wilson and Moehlis, 2015; Yuan *et al*, 2022). Thalamic stimulation in epilepsy patients at 15–45 pps produced synchronization of hippocampal local field potentials, whereas higher frequencies above 45 pps promoted desynchronization of hippocampal activity and reduction of pathological epileptic discharges (Yu *et al*, 2018). Adenosine release could be another factor mediating seizure suppression at higher frequencies (Boison, 2013; Crosson and Gray, 1997; Kakiuchi *et al*, 1969; Loscher and Kohling, 2010; Toprani *et al*, 2013).

It is also notable that the optimal frequencies for seizure induction suggested by our results correspond well with EEG frequencies observed during the onset of epileptic seizures. Gastaut and Broughton described a 10 Hz "recruiting rhythm" in the EEG during the first 10 seconds of generalized tonic-clonic seizures, followed by a progressive decrease in frequency until seizure termination (Gastaut and Broughton, 1972). There has been little investigation of the mechanisms underlying the recruiting rhythm, even in animal models (Kohsaka *et al*, 2002). In vitro studies have shown bursting patterns of ~ 10 Hz in groups of neurons in response to epileptogenic stimuli, which may result from sustained dendritic depolarization (Kawaguchi, 2001; Traub *et al*, 1996). Gastaut and Broughton likened the rhythm to that seen in thalamically-driven cortical responses (Gastaut *et al*, 1972); whether thalamocortical interactions play a role in seizure induction by ECT or MST is currently unknown. Perhaps a "resonance" effect may exist whereby fewer stimulus pulses are required to induce seizures at frequencies characteristic of the onset of endogenous epileptic activity.

Finally, our EEG analysis found stimulus frequency affected ictal expression, suggesting shifts in the dynamics of neural inhibitory and excitatory processes with stimulation frequency.

*Implications for clinical ECT and MST*

The present study evaluated the impact of ECT and MST stimulus frequency on ST and seizure characteristics at threshold in nonhuman primates. The effects of stimulus frequency on the characteristics of suprathreshold seizure induction, therapeutic efficacy, and side effects has to be investigated in future studies. Evaluating efficacy would require appropriately designed clinical trials, since animal models of depression have significant limitations (Kritzer *et al*, 2023). Presently, the literature on the impact of stimulus frequency on clinical outcomes of ECT and MST is limited. Studies that considered the effect of stimulus frequency on the efficiency of seizure induction with ECT found that higher frequencies were less efficient (Devanand *et al*, 1998; Girish *et al*, 2003; Peterchev *et al*, 2010; Swartz and Larson, 1989; Weaver *et al*, 1982), consistent with our results. In one ECT study, stimulus frequency had no effect on heart rate or seizure induction with suprathreshold trains with matched number of pulses at 30 Hz (60 pps) and 60 Hz (120 pps) (Swartz and Manly, 2000). Another trial found no differences in seizure durations, ictal cardiovascular responses, and therapeutic outcome between the 50 pps and 200 pps (Kotresh *et al*, 2004). A third study indicated that 40 Hz (80 pps) is more effective than 100 Hz (200 pps), and there were no significant differences in memory testing (Roepke *et al*, 2011).

Stimulus frequency may impact clinical MST outcomes (Daskalakis *et al*, 2020; Kayser and Wagner, 2018). An MST trial by Daskalakis et al. comparing low (25 pps), medium (50 or 60 pps), and high (100 pps) frequency used a titration of the individual ST by incrementing the number of pulses similar to the present study, except that the pulse amplitude was fixed at MSO for all participants (Daskalakis *et al*, 2020). Treatments were then delivered by increasing the number of pulses relative to the ST for effective suprathreshold stimulation (Backhouse *et al*, 2018; Daskalakis *et al*, 2020). Notably, there were substantial differences in the average number of pulses delivered during treatment for each group: 396, 724, and 753, for low, medium, and high frequency, respectively (Backhouse *et al*, 2018), reflecting lower STs for the low frequencies, consistent with our findings. Regarding efficacy, high frequency produced the highest rates of remission of depression symptoms, followed by moderate and low frequency (Daskalakis *et al*, 2020). However, neither response rates nor number of treatments associated with remission were significantly different across the conditions (Daskalakis *et al*, 2020). Since 100 pps MST was administered with nearly twice the number of pulses for 25 pps MST, it is unclear if the superior remission rate was caused by the higher frequency or the larger number of pulses. The fact that response rates at 100 pps and 25 pps were comparable despite the two-fold difference in number of pulses, leaves open the possibility that 25 pps could be as effective or superior if the number of pulses were matched. Further, while seizure duration was comparable across the stimulus frequency groups (Daskalakis *et al*, 2020), consistent with our findings, a common EEG measure of seizure adequacy was better for 25 pps and 50 pps than for 100 pps (Backhouse *et al*, 2018). On the other hand, the time to reorientation was significantly shorter for the high frequency than the medium or low frequency stimulation, and consistency of autobiographical recall was more affected for the medium frequency than low or high frequency, suggesting that higher frequency stimulation could have an advantageous cognitive side effects profile (Daskalakis *et al*, 2020). Interestingly, an exploratory analysis suggested that the low and medium stimulus frequencies may be more effective that the high frequency at reducing suicidality (Weissman *et al*, 2020), potentially pointing to interactions between stimulus frequency and symptom dimensions.

These findings from clinical studies, while limited, hint at interesting possibilities about the impact of stimulation frequency on seizure characteristics, efficacy, and side effects, but more research is needed. Our results on the relationship between the ECT and MST stimulus frequency and ST can inform such work. One practical challenge is testing low frequency stimuli with an adequate number of pulses. In our data, frequencies above 10 pps required stimulation duration below 10 s, which is practical. However, prior findings that for ECT to be effective the number of stimulus pulses has to exceed the ST by several times (Sackeim *et al*, 2000) would imply relatively long stimulus trains that may surpass what commercially available devices can deliver: maximum of 8 s in modern ECT devices (MECTA Corp., 1997; SigmaStim, 2022; Somatics LLC, 2021) and 20 s in MST devices (Daskalakis *et al*, 2020). Therefore, devices capable of extended trains may be necessary to investigate effective low-frequency stimulation, and the safety of longer train duration has to be assessed. Our preliminary results in nonhuman primates indicate that long trains of 25 pps with individually-titrated current amplitude and lasting up to 40 s appear safe based on heart rate, blood pressure, and blood oxygenation level assessment (Peterchev *et al*, 2016).

*Implications for rTMS*

Our data on ST across stimulus frequencies could potentially inform safety considerations for rTMS, where seizures must be avoided. While there is the potential confound of anesthesia, the anesthetic agents used in the present study are common in ECT and are selected to not increase the ST significantly. With regard to rTMS safety, the assumption has been that higher frequencies carry higher seizure risk (Rossi *et al*, 2021; Rossi *et al*, 2009). The decreasing ST as frequency increases in the 5–25 pps range is consistent with the guidelines for rTMS which specify that the number of pulses that can be safely administered without inducing a seizure decreases similarly in the 1–25 pps frequency range (Rossi *et al*, 2009). However, our results suggest that above 25 pps, the rTMS ST may start to increase; this may contribute to the documented safety of theta-burst stimulation which incorporates 50 pps bursts (Rossi *et al*, 2021) and could inform safety considerations for protocols involving very high frequencies (Jung *et al*, 2016; Jung *et al*, 2021).
